# C4 induces pathological synaptic loss by impairing AMPAR trafficking

**DOI:** 10.1101/2023.09.09.556388

**Authors:** Rhushikesh A. Phadke, Ezra Kruzich, Luke A. Fournier, Alison Brack, Mingqi Sha, Ines Picard, Connor Johnson, Dimitri Stroumbakis, Maria Salgado, Charlotte Yeung, Berta Escude Velasco, Yen Yu Liu, Alberto Cruz-Martín

## Abstract

During development, activation of the complement pathway, an extracellular proteolytic cascade, results in microglia-dependent synaptic elimination via complement receptor 3 (CR3). Here, we report that decreased connectivity caused by overexpression of *C4* (C4-OE), a schizophrenia-associated gene, is CR3 independent. Instead, C4-OE triggers GluR1 degradation through an intracellular mechanism involving endosomal trafficking protein SNX27, resulting in pathological synaptic loss. Moreover, the connectivity deficits associated with C4-OE were rescued by increasing levels of SNX27, linking excessive complement activity to an intracellular endolysosomal recycling pathway affecting synapses.

## MAIN

The immune complement pathway is central to neuroimmune interactions, playing a crucial role in synapse elimination and plasticity^1–3^. Disruptions in the complement pathway have been linked to brain diseases that exhibit pathological synaptic loss, such as schizophrenia (SCZ) and Alzheimer’s Disease^4–7^. We and others have previously demonstrated that increasing the levels of complement component 4 (*C4A* or just *C4*), a key SCZ risk gene, led to pathological synaptic loss in medial prefrontal cortex (mPFC) pyramidal neurons and PFC-associated behavioral deficits in mice^8,9^. The current working hypothesis suggests that complement proteins are released into the extracellular space and modify neuronal connectivity by activating complement receptor 3 (CR3) and recruiting microglia to modify synaptic connections^10^.

To determine whether CR3 is necessary for C4-induced synaptic loss, we overexpressed C4 (C4-OE) via *in utero* electroporation (**Fig. 1A**) in transgenic mice lacking this key phagocytic receptor of the complement cascade^2,11^ (CR3KO) (**Fig. 1B**). Our group previously showed in control experiments that we can reliably increase levels of C4 in cortical synapses *in vivo* using IUE without dramatically altering the intrinsic electrophysiological properties of transfected cells or inducing toxicity^8^. Remarkably, we observed a ∼16% reduction in the density of postsynaptic apical dendritic spines of layer (L) 2/3 cortical pyramidal neurons of postnatal (P) 21-23 mice as compared to GFP-controls, suggesting that C4 acts through a CR3 independent, non-canonical mechanism to drive synaptic loss (**Fig. 1C**).

**Figure 1:**
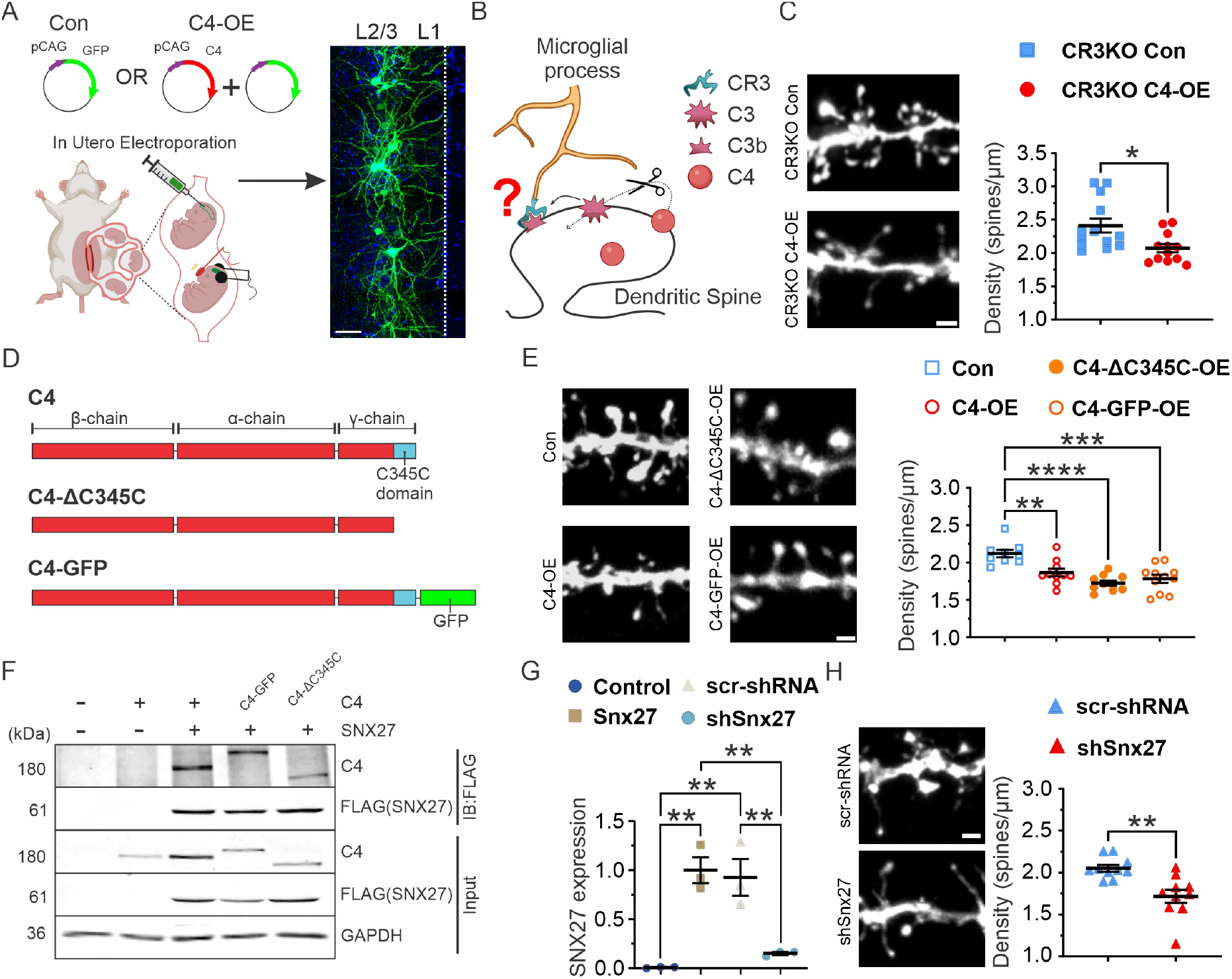
C4 overexpression led to mPFC pyramidal neuron hypoconnectivity through a CR3 independent mechanism. (**A**) Left: *In utero* electroporation (IUE) procedure performed on E16 embryos. GFP-control (Con) represents the transfection of a single plasmid (GFP under the CAG promoter), while C4-OE represents the transfection of two plasmids (GFP and C4, both under the CAG promoter). Right: Representative 60X confocal image of P21-23 L2/3 mPFC neurons transfected with GFP (green). DAPI, cytoarchitecture. White dotted line, pia. Scale bar = 100 μm. (**B**) Model depicting the mechanism of spine removal through C3b recognition by the microglia expressing CR3^4^. (**C**) Left: Representative confocal image (60X) of CR3KO-control (blue squares) and CR3KO C4-OE (red circles). GFP, white signal. Scale bar = 2 μm. Right: C4-OE in mice lacking the CR3 showed decreased dendritic spine density relative to control CR3-KO mice. t test. \**p*<0.05. (**D**) Structure of wt C4 and C4 mutants (C4-ΔC345C and C4-GFP). C4 comprises three main chains (β, α and γ) and a highly conserved C-terminal domain known as C345C (cyan). (**E**) Left: Representative confocal image (60X) of Control and C4 mutants. Right: OE of wt C4 and mutants C4-GFP-OE and ΔC345C-OE caused a decrease in dendritic spine density relative to GFP-control. One-way ANOVA. \*\**p*<0.01, \*\*\**p*<0.001, \*\*\*\**p*<0.0001. Scale bar = 2 μm. (**F**) Western blot (WB) showing co-IP from HEK293T cells transfected with SNX27 and either C4, C4-ΔC345C or C4-GFP plasmids. The presence of C4, SNX27 and GAPDH proteins was detected by Western blotting in HEK293T cell lysates (Input). Interaction of wt C4 and C4 mutants with SNX27 was detected by Western blotting of co-IP against FLAG-tagged SNX27 (IB: FLAG). Protein molecular weights are indicated in kilodaltons (kDa) on the left. + indicated presence of wild type protein. (**G**) HEK293T cells were transfected with either no plasmid (Control, blue circle), Snx27 only (Snx27, yellow squares), Snx27 with scr-shRNA (scrambled control, gray triangles) or Snx27 with Snx27 shRNA (shSnx27, cyan circles). Compared to Snx27 and scr-shRNA control, shSnx27 led to approximately 85% knockdown, normalized to GAPDH loading control. t test. \*\**p*<0.01. (**H**) Left: Representative confocal image (60X) of scr-shRNA control (solid blue triangles) and shSNX27 (solid red triangles). GFP, white signal. Scale bar = 2 μm. Right: SNX27 KD led to a decrease in spine density relative to scr-shRNA. t test. \*\**p*<0.01. (**C**) *N*=13 CR3KO-control dendrites, 3 animals; *N*=12 CR3KO-C4-OE dendrites, 3 animals, (**E**) *N*=9 GFP-control dendrites, *N*=10 C4-OE and C4-ΔC345C-OE dendrites, *N*=11 C4-GFP dendrites, 3 animals each. (**H**) *N*=10 scr-shRNA control and shSNX27 dendrites, 3 animals each. All graphs, Mean ± SEM.

Consistent with C4-OE causing synaptic loss independent of complement activation, overexpression of C4 mutants that lack the enzymatic activity of C3 convertase -an enzyme essential for the generation of CR3 ligand (C3b) - by either the addition of a sterically large GFP tag (C4-GFP) or by deletion of the C345C domain (C4-ΔC345C)^12^ (**Fig. 1D**) resulted in a ∼22% decrease in spine density (**Fig. 1E**), compared to GFP-control mice. Therefore, C4-induced synaptic loss is independent of activation of the downstream complement cascade.

Next, we leveraged the protein interactome (BioPlex) in human cells to identify sorting nexin 27 (SNX27) as a potential interacting partner of C4^13^, which we validated by performing a co-immunoprecipitation (co-IP) assay in HEK293T cells (**Fig. 1F**) using C4 and FLAG-tagged SNX27. SNX27 is an endosomal protein that recycles glutamatergic AMPA receptors (R) to the postsynaptic surface^14–16^, strengthening synapses via long-term potentiation^17^. We also found that C4-GFP and C4-ΔC345C both bind to SNX27, suggesting that C4 mutants with disrupted C3 convertase activity do not interfere with C4-SNX27 interactions (**Fig. 1F**).

To determine whether C4 and SNX27 were likely to interact in neurons, we used STED nanoscopy in dissociated mouse mPFC neuronal cultures (DIV 21) overexpressing *C4* and FLAG-tagged *Snx27*. In dendrites of cultured neurons, we identified instances of C4 colocalized with SNX27 (**Extended Data Fig. 1**), suggesting that these proteins are in proximity to one another within the dendrites. Additionally, knockdown (KD) of endogenous Snx27 in mPFC neurons with shRNA against SNX27 (**Fig. 1G**) resulted in a ∼20% decrease in spine density as compared to the scrambled (scr)-shRNA control (**Fig.1H**), mimicking the C4-OE synaptic phenotype that we saw *in vivo*.

Thus, our findings challenge established views on the mechanism by which complement pathway’s role in synaptic plasticity by showing that high levels of C4 reduce synaptic connectivity independent of CR3. This is supported by decreased connectivity seen following overexpression of C4 mutants that lack C3 convertase activity, co-IP of C4 with SNX27, colocalization of C4 and SNX27 in dendrites, and Snx27 KD phenocopying the C4-OE-induced effects, indicating a novel neuronal pathway in which C4 can modify synaptic pathologies.

By interactions through its PDZ domain, SNX27 plays a pivotal role in boosting the recycling of glutamate AMPARs and NMDARs from early endosomes to the plasma membrane^14–16^, safeguarding postsynaptic receptors from degradation and promoting synapse formation^18^. To determine whether increased expression of SNX27 in the C4-OE background restored spine deficits, we simultaneously overexpressed C4 and SNX27 (C4-SNX27-OE) (**Fig. 2**). Dendritic spine density in C4-SNX27-OE was indistinguishable from GFP-control, rescuing the C4-induced phenotype (**Fig. 2A, B**) and suggesting that upregulation of this endosomal protein can counteract C4-OE-driven hypoconnectivity. Notably, overexpression of SNX27 alone (SNX27-OE) did not increase spine density compared to GFP-control (**Fig. 2A, B**), demonstrating that the SNX27 rescue drives structural spine plasticity through a C4-specific cellular mechanism rather than a general increase in structural synaptic plasticity.

**Figure 2:**
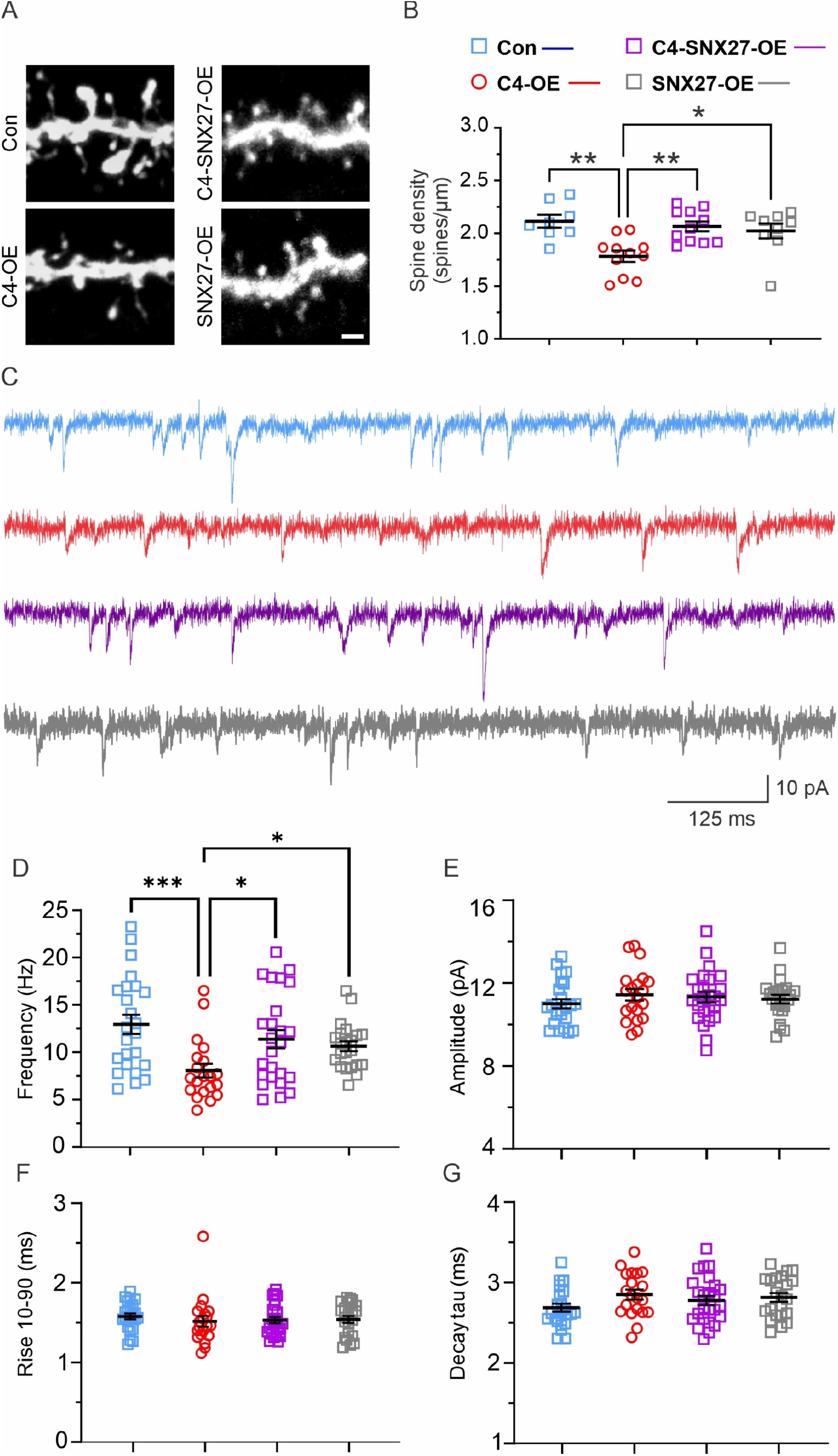
Increasing neuronal levels of SNX27 rescued the C4 hypoconnectivity phenotype. (**A**) Representative confocal image (60X) of GFP-positive P21-23 L1 apical tufts in GFP-control (Con, blue squares), C4-OE (red circles), C4-SNX27-OE (purple squares) and SNX27-OE (gray square) in the mPFC. GFP, white signal. Scale bar = 2 μm. (**B**) C4-OE led to decreased dendritic spine density relative to GFP-control. However, this synaptic pathology was rescued by simultaneous OE of SNX27 (C4-SNX27-OE). SNX27-OE on its own did not lead to changes in dendritic spine density. One-way ANOVA. \**p*<0.05, \*\**p*<0.01. (**C**) Representative traces of mEPSCs in GFP-control (blue), C4-OE (red), C4-SNX27-OE, (purple), and SNX27-OE (gray) L2/3 mPFC pyramidal neurons from P20-25 animals. Scale bars, 125 ms,10 pA. (**D**) mEPSC frequency was decreased in C4-OE relative to GFP-control (Kruskal-Wallis one-way ANOVA. \*\**p*<0.01; GFP-control vs. C4-OE, Dunn’s multiple comparison test. \*\*\**p*<0.001) but was rescued when C4 and SNX27 were co-overexpressed (C4-OE vs. C4-SNX27-OE, Dunn’s multiple comparison test. \**p*<0.05). SNX27-OE alone did not alter mEPSC frequency relative to GFP-control (Con vs. SNX27-OE, Dunn’s multiple comparison test. *p*>0.9999) but was significantly greater than in C4-OE (C4-OE vs. SNX27-OE, Dunn’s multiple comparison test. \**p*<0.05). Hz; Hertz. (**E-G**) There were no differences in the amplitude (**E**, One-way ANOVA. *p*=0.6241), Rise 10-90 (**F**, Kruskal-Wallis One-way ANOVA. *p*=0.4346), and Decay tau (**G**, One-way ANOVA. *p*=0.2122). *N*=24 GFP-control cells, from 6 animals. *N*=20 C4-OE cells, from 4 animals. (**B**) *N*=8 control dendrites, N=9 C4-SNX27-OE dendrites, N=11 C4-OE and SNX27-OE dendrites, 3 animals each. (**D-G**) *N*=25 C4-SNX27-OE cells, from 4 animals. *N*=22 SNX27-OE cells, from 5 animals. All graphs, Mean ± SEM.

Next, we determined whether the C4-OE induced spine loss and subsequent rescue of these deficits with C4-SNX27-OE (**Fig. 1E** and **Fig. 2A, B**) resulted in parallel changes at a functional level (**Fig. 2C-G**). To test this, we performed electrophysiological whole-cell voltage-clamp recordings in acute brain slices prepared from the mPFC to monitor the excitatory drive of transfected cortical neurons. We found that increased levels of C4 led to a ∼38% reduction in the frequency of miniature excitatory postsynaptic currents (mEPSCs) (**Fig. 2C, D**) as compared to GFP-control, indicating that C4-OE causes a decrease in functional synaptic connectivity. We also found that C4-SNX27-OE mice did not show this functional deficit, restoring mEPSC frequency to GFP-control levels **(Fig. 2D**). Importantly, SNX27-OE did not alter mEPSC frequency relative to controls (**Fig. 2D**), indicating that SNX27 rescues the hypoconnectivity phenotype through a C4-specific mechanism. In contrast to our previous study^8^, we did not identify any significant changes in mEPSC amplitude between groups (**Fig. 2E**). One reason for the discrepancy might be the different recording solutions used in the two studies. Consistent with our previous study, we did not observe any significant differences in mEPSC rise or decay kinetics (**Fig. 2F** and **Fig. 2G**, respectively).

These findings indicate that increasing levels of the endosomal protein SNX27 rescues the C4-OE-induced hypoconnectivity phenotype, suggesting a possible link between overexpression of C4, disruption of the synaptic trafficking machinery, and pathological synaptic loss.

SNX27 mediates the insertion of AMPAR subunits into the postsynaptic membrane^14^, which is necessary to stably increase spine size^17^ and promote the effects of experience on synaptic transmission efficacy *in vivo*^18,19^. In the past, we demonstrated that C4-OE impacts the circuitry of the L1 apical tufts^8^. Axonal projections to superficial L1 are part of a circuit that controls consciousness, attention, and learning states^20^. Therefore, we sought to determine whether C4-OE alters the recycling of AMPA GluR1 subunits in the L1 dendritic spine apical tufts of transfected mPFC pyramidal cells (**Fig. 3**).

**Figure 3:**
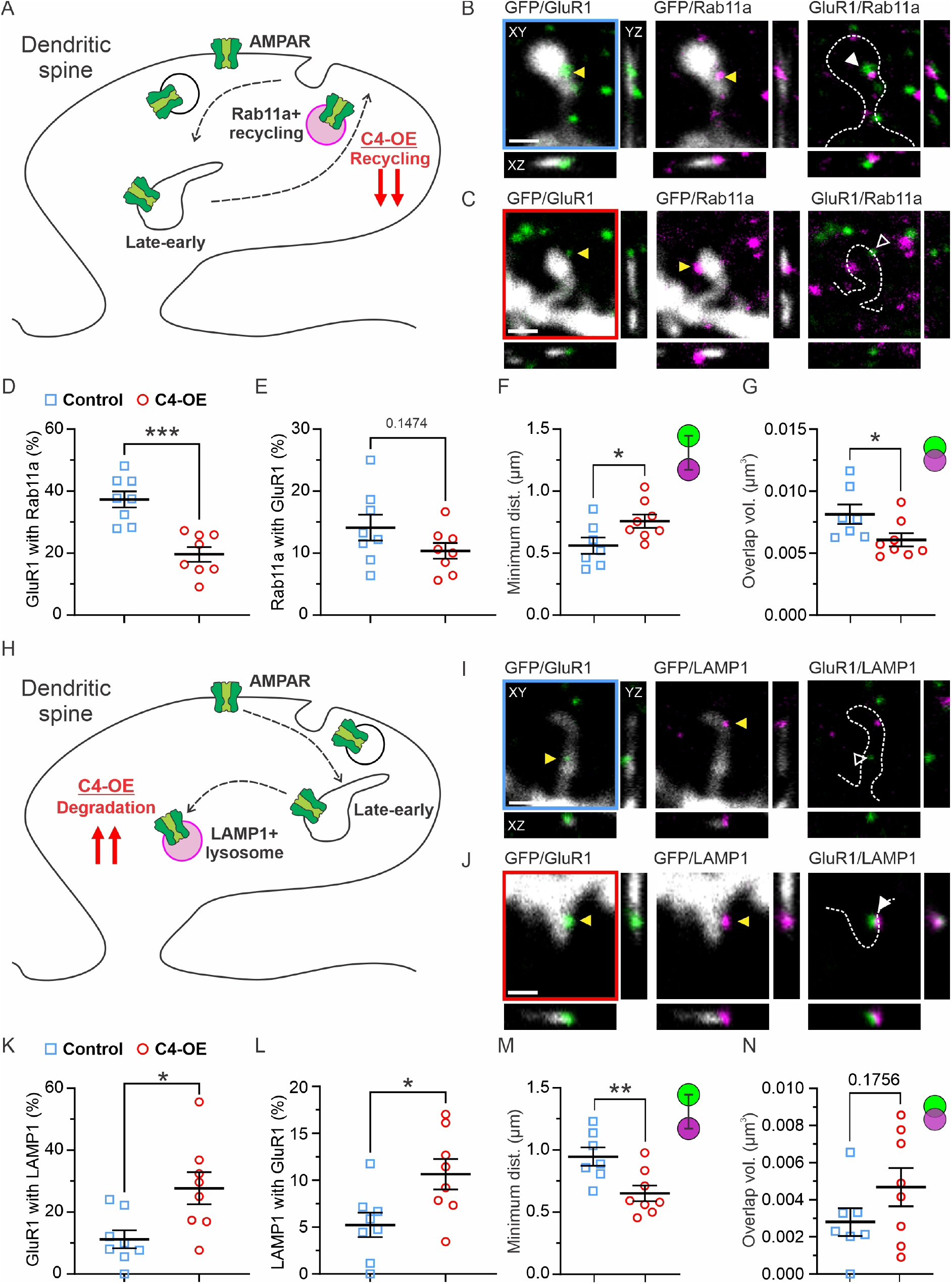
Super-resolution imaging revealed that C4-OE led to decreased recycling and increased degradation of GluR1-containing AMPAR in L1 apical tuft dendritic spines. (**A**) Model depicting effects of C4 overexpression on GluR1 recycling in dendritic spines. (**B**) Representative images (60X) showing a GFP-positive dendritic spine (white), GluR1 (green) and Rab11a (magenta) of P21-23 apical tufts in GFP-controls (blue frame). Orthogonal views are shown (XY, YZ and XZ). Yellow arrowhead indicates an instance of GluR1 or Rab11a clusters in spines. White-filled arrowhead indicates an instance of GluR1/Rab11a colocalization. Spine silhouette, white dotted line. Scale bar = 2 μm. (**C**) Representative images (60X) showing a dendritic spine identified with GFP signal (white), GluR1 (green) and Rab11a (magenta) in C4-OE (red frame). Orthogonal views are shown (XY, YZ and XZ). Yellow arrowhead indicates an instance of GluR1 or Rab11a clusters in spines. White empty arrowhead indicates an instance of non-colocalized GluR1/Rab11a. Spine silhouette, white dotted line. Scale bar = 2 μm. (**D**) C4-OE caused a 47% decrease in the amount of GluR1 colocalized with Rab11a compared to GFP-control. (**E**) There was no change in the amount of Rab11a colocalized with GluR1 in C4-OE relative to GFP-control. (**F**) C4-OE increased the minimum distance between GluR1 and Rab11a clusters by 35% relative to GFP-control. Green circle: GluR1, Magenta circle: Rab11a. (**G**) C4-OE led to a 25% decrease in the overlapping volume between GluR1 and Rab11a relative to GFP-control. Green circle: GluR1, Magenta circle: Rab11a. (**H**) Schematic showing the effects of C4-OE on GluR1 degradation in dendritic spines. (**I**) Representative images (60X) showing a GFP-positive spine (white), GluR1 (green) and LAMP1 (magenta) in GFP-control (blue frame). Orthogonal views are shown (XY, YZ and XZ). Yellow arrowheads indicate instances of GluR1 or LAMP1 clusters present in spines. White empty arrowhead indicates an instance of non-colocalized GluR1/LAMP1. Spine silhouette, white dotted line. Scale bar = 2 μm. (**J**) Representative images (60X) showing a GFP-positive spine (white), GluR1 (green) and LAMP1 (magenta) in C4-OE (red frame). Orthogonal views are shown (XY, YZ and XZ). Yellow arrowhead indicates an instance of GluR1 or Rab11a clusters in spines. White-filled arrowhead indicates an instance of GluR1/LAMP1 colocalization. Spine silhouette, white dotted line. Scale bar = 2 μm. (**K**) C4-OE led to a 145% increase in the amount of GluR1 colocalized with LAMP1. (**L**) C4-OE caused a 103% increase in the amount of LAMP1 colocalized with GluR1. (**M**) C4-OE induced a 31% decrease in the minimum distance between GluR1 and LAMP1 clusters. Green circle: GluR1, Magenta circle: LAMP1. (**N**) Compared to GFP-control, C4-OE did not alter the overlapping volume between GluR1 and LAMP1. Green circle: GluR1, Magenta circle: LAMP1. (**D-E, K-L**) *N*=8 dendrites, 3 animals for Con and C4-OE. (**F-G, M-N**) *N*=7 dendrites, 3 animals for Con and *N*=8 dendrites, 3 animals C4-OE. (**D-G, K-N**) t test. \**p*<0.05, \*\**p*<0.01, \*\*\**p*<0.001. All graphs, Mean ± SEM.

To do this, we first used immunohistochemistry to stain for GluR1 and the recycling endosome protein Rab11a^21^, followed by STED nanoscopy in fixed mPFC tissue sections (**Fig 3A-C**). Rab11a triggers the cellular machinery required to sort and recycle membranes and proteins back to the plasma membrane. STED imaging allowed us to inspect the nanoscale organization of the spine by identifying colocalized (**Fig. 3B**) and non-colocalized (**Fig. 3C**) GluR1 and Rab11a protein clusters in spine subcompartments. Additionally, we found that C4-OE caused a decrease in the quantity of GluR1 clusters colocalizing with Rab11a, while the quantity of Rab11a-positive clusters associating with GluR1 showed a decreasing trend without reaching significance (**Fig. 3D, E**). Furthermore, C4-OE in cortical neurons also caused an increase in the minimum distance (**Fig. 3F**) and a decrease in the degree of overlap (**Fig. 3G**) of colocalized GluR1-Rab11a clusters, suggesting that C4-OE decreased the association of GluR1 with Rab11a-positive recycling endosome, disrupting the recycling of the receptors to the plasma membrane.

To determine whether C4-OE triggers degradation of GluR1, we next imaged GluR1 and LAMP1-positive lysosomal clusters^22^ (**Fig. 3H-J**). LAMP1 is a glycoprotein that is a major component of the lysosomal membrane. We found colocalized (**Fig. 3I**) and non-colocalized (**Fig. 3J**) GluR1 and LAMP1 clusters in spine compartments. We observed that C4-OE caused increased instances of GluR1 colocalizing with LAMP1 and increased LAMP1 associating with GluR1 (**Fig. 3K, L**). Additionally, we observed a decrease in the minimum distance between GluR1 and LAMP1 clusters (**Fig. 3M**), with no significant change in their degree of overlap with increased levels of C4 (**Fig. 3N**). These results suggest that C4-OE increased the association of GluR1 with the lysosomal degradation machinery. Compared to GFP-control, increased levels of C4 did not alter the size of GluR1, Rab11a, and LAMP1 clusters in the spine (**Extended Data Fig. 2**). Overall, these findings demonstrate that elevated levels of neuronal C4 disrupt GluR1 recycling, favoring GluR1 degradation instead, thus contributing to impaired synaptic plasticity.

We have demonstrated that the decreased connectivity caused by increased expression of *C4*, a gene associated with SCZ, is not dependent on CR3, the main phagocytic receptor that drives complement-mediated synaptic engulfment. Instead, using STED nanoscopy and electrophysiological approaches, we showed that C4-OE leads to the degradation of GluR1, which possibly causes the loss of synapses in the mPFC, a region associated with devastating neuropsychiatric disorders^23^. This is further supported by increased levels of SNX27 rescuing the synaptic phenotype in *C4* overexpressing cortical neurons, which suggests that excessive complement activity can affect synaptic plasticity through the dysregulation of GluR1-containing AMPA receptor recycling to the postsynaptic membrane.

Therefore, we propose a new model where increased neuronal levels of C4 lead to deficits in GluR1-containing AMPA receptor recycling to postsynaptic membranes and an increased degradation of GluR1 in the spine lysosome^14,16^. Future research will decipher whether interactions between C4 and SNX27 are direct or occur through a larger protein complex in the postsynaptic spines, as well as whether this interaction is specific to GluR1 or impacts the trafficking of other glutamate receptor subunits similarly.

An alternate explanation of the results is that increasing the levels of SNX27 leads to C4 sequestration and limits its availability, which would potentially block microglia-dependent synaptic pruning and inhibit synaptic loss. We believe that this is highly unlikely because we observed C4-induced synaptic loss in the CR3KO background and by overexpression of C4 mutants that lack C3 convertase activity. In addition, we showed that increased levels of C4 on its own lead to increased degradation of GluR1-containing AMPAR subunits, an established mechanism of synaptic plasticity^18,19,24^ that is not dependent on microglia-mediated synaptic engulfment. These points together indicate that synaptic pruning is not dependent on canonical complement pathways. Lastly, the commonly proposed mechanism of C4-OE-driven synaptic loss requires enzymatic activity in the extracellular space, however SNX27 is an intracellular protein that would not interfere with these C4 properties to limit synaptic engulfment. Even if SNX27 sequestered some C4 protein, a small amount of C4 could be sufficient to drive the local pruning of synapses by microglia due to the amplification mechanisms of the complement pathway. However, that is contradictory to our observation, as SNX27 completely rescues the C4 synaptic phenotype when both proteins are overexpressed.

Previous research has shown that complement overactivation leads to the increased accumulation of synaptic material in the microglia lysosome^2–4,8,10,25^, which has been interpreted as enhanced microglia-mediated synaptic phagocytosis. We postulate that microglia could still play a role in synaptic phagocytosis, but that they do this by eliminating already weakened synaptic connections or their molecular remnants. In this scenario, microglia-dependent synaptic phagocytosis could be triggered by disrupted synapse formation or an inability to stabilize the synapse. In support of this, our group, as well as others, have demonstrated that C4-OE can result in specific deficiencies in filopodia, the precursors of spine synapses, and a decrease in the proportion of newly acquired dendritic spines^8,9^. This indicates that rather than accelerating synaptic pruning, increased levels of C4 may contribute to pathological synaptic loss by interfering with the creation and growth of synapses, which are linked to SNX27 function, glutamate receptor recycling, and long-term potentiation^24^. Nevertheless, our data suggests that CR3-dependent microglial activity alone cannot fully explain the C4-OE-induced synaptic pathology.

Our findings reveal a new mechanism of C4-mediated synapse removal. This discovery is particularly significant in comprehending the wiring of synapses and diseases that stem from complement-driven synaptic loss. Since treatments that modify the complement pathway or focus on microglia-mediated pathological pruning could have negative impacts on the nervous and immune systems, our findings provide a fresh possibility to address the effects of pathological complement activity on the synapse.

## FUNDING AND ACKNOWLEDGEMENTS

This work was supported by a National Institutes of Health R01 (NIMH, 1R01MH129732-01) to A.C-M.; and a Brenton R. Lutz Award to R.A.P. We thank Drs. Thomas Gilmore, J. Tiago Gonçalves, Angela Ho, Ashley L. Comer, and members of the Cruz-Martín lab for critical reading of the manuscript and helpful discussions. We would like to thank Dr. Michael Kirber and the Cellular Imaging Core for providing support with STED experiments and Dr. Todd Blute and the Boston University Biology Imaging Core for providing support for the confocal microscope,

## AUTHOR CONTRIBUTIONS

R.A.P. and A.C-M. conceptualized experiments including formulating composition, goals, and scope of the paper and approaches for analyses. R.A.P., E.K., L.A.F., A.B., M.S., I.P., C.J., C.Y., B.E.V. and Y.Y.L. collected the data and performed experiments. R.A.P., E.K. and L.A.F. performed data curation. R.A.P., E.K., L.A.F., A.B., I.P. and D.S. analyzed data. R.A.P. and L.A.F. contributed code for data analysis. R.A.P., A.C-M., E.K., L.A.F., A.B. and M.S. contributed to parts of the original draft, including figure design and generation. All authors contributed to revision and editing of the draft. A.C-M. obtained funding and supervised the project providing mentorship, oversight, and project administration.

## METHODS

### DNA constructs

For GFP-control, we used a plasmid containing EGFP under the CAG promoter. pCAG-GFP^26^ was a gift from Connie Cepko (Addgene plasmid # 11150 ; http://n2t.net/addgene:11150 ; RRID:Addgene_11150). For C4-OE, DNA sequence containing mouse *C4b* (NM_009780.2, synthesized by Genescript) was subcloned (InFusion Kit, Clonetech) into the pCAG backbone to produce pCAG-C4^8^. C4-GFP was made by inserting GFP from the pCAG-GFP plasmid C terminus to C4 using a GSSGSS linker (subcloned by Genescript). For co-electroporation with C4-GFP, mRuby3 under pCAG promoter was used. pCAG-mRuby3-WPRE was a gift from Rylan Larsen (Addgene plasmid # 107744 ; http://n2t.net/addgene:107744 ; RRID:Addgene_107744). C4-ΔC345C was made by deleting amino acids 1589-1736 using plasmid amplification. For SNX27-OE, we used a plasmid containing SNX27-FLAG under CMV promoter (pCMV-SNX27-FLAG, OriGene Technologies plasmid #MR218832). For SNX27 knockdown experiments, we used plasmids containing scrambled shRNA and anti-SNX27 shRNA under the CMV promoter (pCMV-scr-shRNA, pCMV-shSNX27, Origene Technologies #TL518223). All plasmid DNA were purified using the ZymoPure II (Zymo Research) plasmid preparation kit and were resuspended in molecular biology grade water.

### HEK293T cell culture

#### Preparation and maintenance

HEK293T cells were maintained in a 10 cm tissue culture dish (Genesee Scientific, 25-202) in 10 ml of complete media: DMEM (Cytiva, SH30243.01) supplemented with 10% fetal bovine serum (Cytiva, SH30396.03), penicillin (100 units/ml), and streptomycin (100 μg/ml) (Sigma-Aldrich, P4458) following protocols described previously^8^. Cells were kept at 37°C and 5% CO_2_ in a Thermo Scientific HERAcell 150i incubator. Cells were passaged until they reached 90-95% confluency and seeded into a new 10 cm dish at a seeding density of 1 x 10^6^ for maintenance.

#### Transfection

HEK293T cells were seeded in a 6-well plate (Genesee Scientific, 25-105) at a density of 0.2 x 10^6^ in 2 ml of complete media as described above. Two days after seeding, the cells were transfected using Geneglide transfection reagent (Biovision, M1081-1000). The complete media in the 6-well plate was replaced by 1 ml per well of DMEM. Two sets of tubes were prepared per transfection condition. Tube 1 contained 500 μl of DMEM along with Geneglide reagent, and Tube 2 contained 500 μl of DMEM along with plasmid DNA for transfection. One μg of each plasmid was used per well, and 2 μl of Geneglide per μg of plasmid was used per well. The contents of Tube 2 were mixed with a pipette and transferred gently to Tube 1. After invert mixing the contents, they were incubated at room temperature for 20 min, and added drop by drop to the 6-well plate, followed by gentle swirling. The plate was incubated for 24 h at 37 °C and 5% CO_2_ after which the transfection media was replaced by 2 ml of complete media.

### Co-immunoprecipitation and Western blotting

#### Co-immunoprecipitation

HEK293T cells were transfected with three possible conditions: Control with no plasmid, only C4 plasmid, and multiple combinations of two plasmids for testing protein interactions, for example wt SNX27 and C4 mutant plasmids. 48 h after transfection (as described above), the 6-well plate was removed from the incubator and put on ice. The media in each well was removed, cells were washed with PBS and lysed with 500 μl of lysis buffer consisting of 0.5% NP40 (Boston BioProducts, P-877-250), protease inhibitor cocktail 1X (Thermo Fisher, 87786), 30 mM Tris and 150 mM NaCl, pH 7.5 in Milli-Q water. After incubation on ice for 15 min, the cells were collected using a cell scraper and transferred to a 1.7 ml microcentrifuge tube. The tube was sealed and rocked with end-over rotation overnight at 4 °C. Following rotation, the tubes were centrifuged at 13,000 rpm for 30 min, and the supernatant was collected for further analysis as a cell lysate. Next, 200 μl of the lysate was incubated with 1:100 dilution of rabbit anti-FLAG antibody (Abcam, ab205606) overnight at 4 °C with end-over rotation. The lysate-antibody mix was then incubated with Magnetic Protein A beads (New England Biolabs, S1425S) overnight at 4 °C with end over rotation. Following incubation with magnetic beads, the beads were separated from the liquid using a neodymium magnet and washed 3 times with 100 μl buffer consisting of 30 mM Tris and 150 mM NaCl in Milli-Q water. Finally, 30 μl of SDS loading buffer (BioRad, 1610747 with 10% 2-Mercaptoethanol) was added to the beads and they were incubated at 95 °C for 5 min to elute protein complexes before using them for western blots.

#### Western blotting

HEK293T cell lysate and co-immunoprecipitation elutes were obtained as described above. Samples were size-separated using SDS-PAGE with a 4-20% gradient gel, followed by transfer onto a PVDF membrane (Cytiva, 10600021) for Western blotting analysis. The transfer was carried out at 100 mA constant current, overnight at 4 °C, using a transfer buffer containing 20% methanol, 25 mM Tris pH 7.0 and 100-mM Glycine. Following transfer, the membrane was incubated with a blocking buffer containing 5% nonfat dry milk powder, 0.1% Tween-20, and 0.1% sodium azide in 1X Tris-buffered saline (TBS) (Genesee Scientific, 18-236B), for 1 h at room temperature with tilt shaking. The blocking buffer was removed and replaced with a primary antibody solution (diluted in blocking buffer). The membranes were incubated with primary antibodies overnight at 4 °C with tilt shaking. After incubation, the primary antibody solution was removed and the membranes were washed three times for 10 min each at room temperature with TBST: 0.1% Tween-20, 1X TBS. The membrane was then incubated with a secondary antibody solution (diluted in blocking buffer) overnight at 4 °C with tilt shaking, followed by three washes with TBST. The membrane was then stored in TBST until imaging.

#### Analysis of Western blots

Western blot membranes were imaged using Sapphire Biomolecular Imager (Azure Biosystems, Dublin, CA). Depending on the choice of secondary antibody, either 488 or 640 nm wavelengths were used for the image acquisition. The acquired images were then analyzed in ImageJ using the Gel Analysis Plugin to get intensity values for bands. Each gel run was accompanied by a well containing a protein ladder (Thermo Fisher Scientific, PI26625, Waltham, MA) to help identify the correct molecular weights of bands stained by antibodies. In cases of expression analysis between different conditions, GAPDH (Glyceraldehyde 3-phosphate dehydrogenase) was used as a normalization control.

#### Antibodies used for Western blotting

The following primary antibodies were used for staining: Rabbit anti-C4 (1:300, Biogen), Mouse anti-GAPDH (1:1000, Santa Cruz Biotechnology, sc-47724), Mouse anti-FLAG (1:1000, Sigma-Aldrich, F3165). The following secondary antibodies were used for staining: Goat anti-rabbit 647 (1:1000, Sigma-Aldrich, 40839-1ML-F), Goat anti-mouse 488 (1:1000, Jackson ImmunoResearch, 115-545-003).

## Ethics statement

All experimental protocols were conducted according to the National Institutes of Health (NIH) guidelines for animal research and were approved by the Boston University Institutional Animal Care and Use Committee (IACUC; protocol #2018-539).

## Animals

All mice were group housed on a 12-h light and dark cycle with the lights on at 7 AM and off at 7 PM and with food and water *ad libitum*. Offspring were housed with dams until weaning at P21. Experiments were performed in CD-1 wild-type mice of both sexes (CD-1 IGS; Charles River; strain code: 022). P21-23 mice were used for STED and spine density experiments. P20-25 mice were used for electrophysiology experiments. For a subset of spine density experiments we used P21-23 CR3-KO mice in a C57BL/6 genetic background (The Jackson Laboratory, B6.129S4-*Itgamtm1Myd*/J, Strain #:003991, RRID:IMSR_JAX:003991^11^).

### *In utero* electroporation

L2/3 progenitor cells in the mPFC were transfected via IUE^8^. Unless otherwise stated, IUE experiments were performed in two-to three-month-old CD-1 dams. All plasmids were used at a final concentration of 1 μg/μl. To generate GFP-control, we electroporated with the pCAG-GFP plasmid. To generate C4-OE, we co-electroporated the pCAG-GFP and pCAG-C4 plasmids. To overexpress C4 mutants, we co-electroporated the pCAG-GFP plasmid with pCAG-C4-ΔC345C plasmid to generate C4-ΔC345C-OE and we electroporated pCAG-mRuby3 with pCAG-C4-GFP plasmid to generate C4-GFP-OE.

To generate SNX27-OE, we co-electroporated the pCAG-GFP with the pCMV-SNX27-FLAG; and to generate C4-SNX27-OE, we co-electroporated the pCAG-GFP, pCAG-C4 and pCMV-SNX27-FLAG. For SNX27 knockdown experiments, pCAG-EGFP was co-electroporated with the pCMV-scr-shRNA or pCMV-shSNX27 for scr-shRNA and shSNX27 groups, respectively. For experiments using the CR3KO mice, we electroporated the pCAG-GFP plasmid in the CR3KO mouse to generate CR3KO-control and co-electroporated the pCAG-EGFP and pCAG-C4 plasmid to generate CR3KO-C4-OE.

Prior to surgery, all tools were sterilized by autoclaving. Aseptic techniques were maintained throughout the procedure, and a sterile field was prepared before surgery using sterile cloth drapes. Animals were weighed, and a combination of buprenorphine (3.25 mg/kg; SC, Bimeda, Inc) and meloxicam (1-5 mg/kg; SC, Covetrus North America) was administered as a preoperative analgesic. Timed-pregnant female CD-1 or CR3KO mice at E16 were anesthetized by inhalation of 4% isoflurane (Covetrus North America) and maintained with 1-1.5% isoflurane via mask inhalation. The abdomen was sterilized with 10% povidone-iodine and 70% isopropyl alcohol (repeated 3 times) before a vertical incision was made in the skin and then in the abdominal wall. The uterine horn was then exposed to allow injection of 0.5-1.0 μl of DNA solution (containing 1 μg/μl plasmid and 0.1% Fast Green) into the lateral ventricles using a pressure-injector (Picospritzer III, Parker Hannifin) with pulled-glass pipettes (Sutter Instrument, BF150-117-10). To target L2/3 progenitor cells in PFC for imaging and electrophysiological experiments, a custom-built triple electrode probe was placed by the head of the embryo^27^, with the negative electrodes placed near the lateral ventricles and the positive electrode placed just rostral of the developing PFC. Next, four-square pulses (pulse duration: 50 ms, pulse amplitude: 36 V, interpulse interval: 500 ms) were delivered to the head of the embryo using a custom-built electroporator. Embryos were regularly moistened with warmed sterile PBS during the surgical procedure. After electroporation, the embryos and uterine horn were gently placed back in the dam’s abdominal cavity and the muscle and skin were sutured (using absorbable and nonabsorbable sutures, respectively). Finally, the dams were allowed to recover in a warm chamber for 1 h and then returned to their cage.

### Primary neuronal culture

Cortical primary neurons were isolated and cultured according to Beaudoin et al.^28^, with minor adjustments. Briefly, E18 pregnant CD-1 dams were deeply anesthetized with an overdose of isoflurane, and an abdominal incision was made to access the uterine horn. Embryos were removed and moved to ice-cold dissociation media as described in Beaudoin et al.^28^. After extraction from the skull, the mPFC was dissected from each hemisphere and collected in fresh dissociation media. After dissection, the tissue was washed and digested with trypsin (Worthington, # LS003707) solution for 20 min at 37 °C, and then Benzonase (Sigma-Aldrich, #E1014-5KU) was added for an additional 5 min. Digested tissue was washed gently twice with dissection media, then twice with plating media, before being resuspended in plating media and gently triturated with 10 and 1 ml pipettes. Cells were plated on poly-L-lysine coated coverslips at a density of 250,000 cells, and the media was aspirated and changed to maintenance media after allowing the cells to adhere for 6 h. After 48 h, cytosine arabinoside (araC, Sigma-Aldrich C1768) was added to a final concentration of 4 μM to inhibit non-neuronal cell growth, and then media was changed again after another 24 h to maintenance media without araC. Neurons were grown for 14 days before fixation with 4% paraformaldehyde + 4% glucose before antibody staining.

### STED super-resolution microscopy

#### Perfusions and Immunohistochemistry

Mice were administered an overdose of sodium pentobarbital (250 mg/kg, Patterson Veterinary, 07-894-1075) intraperitoneally before transcardial perfusion with phosphate buffered saline (PBS, Gibco, Life Science Technologies, 70011044) then 4% paraformaldehyde (PFA) in PBS. The brains were extracted and postfixed in 4% PFA for 24 h, then transferred to a 30% (w/v) sucrose solution and stored at 4 °C. The tissue was sectioned at 40 um thickness with a freezing stage sliding microtome (Leica Biosystems, SM2000). To prepare for immunostaining, the slices were blocked and permeabilized in a blocking buffer: 10% donkey serum (Sigma-Aldrich, S30-100ML) + 1% Tritonx100 (Sigma-Aldrich, X100-100ML) in PBS. Next, the brain slices were incubated with primary antibodies on a shaker at 4 °C for 48 h and then with secondary antibodies in the same fashion. After primary incubation, slices were rinsed with 0.025% Tritonx100 in PBS for 15 min, four times. After secondary incubation, slices were rinsed with PBS for 15 min, four times. The brain slices were mounted onto 1 mm microscope slides (Corning, Cat #:2975-225, Corning) with DAPI-free mounting media (Thermo Fisher Scientific, p36965).

Depending on the experiment, the following primary antibodies were used: Rabbit anti-C4 (1:500, Biogen), Mouse anti-FLAG (1:1000, Sigma-Aldrich, F3165), Mouse anti-GluR1 (1:250, NBP2-22399, Novus Biologicals), Rabbit anti-Rab11a (1:250, Fisher Scientific, 71-5300), Rabbit anti-LAMP1 (1:500, abcam, ab24170). The following secondary antibodies were used: anti-Mouse Abberior STAR Orange (1:300, Fisher Scientific, NC1933863), and anti-Rabbit Abberior STAR Red (1:1000, Fisher Scientific, NC1933870).

#### Imaging

Fluorescence data was acquired using an inverted laser scanning confocal microscope (Nikon Instruments, Nikon Ti2 with Perfect Focus, Plan apochromat 60x 1.4 NA oil immersion objective, Physik Instrumente P736 Piezo Z-stage) outfitted with an Abberior Facility Line STED system controlled with the Abberior Imspector software platform. The 485 nm (for GFP-cell fill), 561 nm (Abberior STAR Orange), and 640 nm (Abberior STAR Red) pulsed excitation lasers and a 775 nm STED depletion laser (Abberior STAR Orange Abberior STAR Red) were used to obtain 25 nm pixel size. For dendrite, synaptic marker, and endosomal marker imaging, ROIs localized to apical dendritic tufts in L1 were acquired consisting of stacks of images (first 50 μm below the pia, approximately 20-40 optical sections, Z-step = 0.2 μm) and saved in OBF file format.

#### Image processing and deconvolution

OBF image files were converted to 32-bit TIFF format and combined to make a single 3-channel image in ImageJ, then imported to IMARIS and converted to IMS file format. To compensate for the difference in resolving power on the Z axis, Z-axial distances were adjusted from 0.2 to 0.1 μm before deconvolution^29^. Using the IMARIS image processing module, the channels for synaptic and endosomal markers (Abberior STAR Orange, Abberior STAR Red) were deconvolved with 10 iterations of the Robust (iterative) algorithm, with Z axial denoising of filter size 0.7.

#### IMARIS Reconstruction

Dendrites were digitally reconstructed in IMARIS using the software’s semi-automatic filament reconstruction function, based on the GFP signal. Dendrite shafts were aligned using seedpoints and spines were reconstructed by hand using the create function and auto-align feature. The Approximate Circle of Cross Section Area algorithm was used for the calculation of both the dendrite and spine diameters. To increase the confidence in puncta reconstruction for analysis, each punctate signal was reconstructed as spots and surfaces. Spots and surfaces were used to describe protein clusters and the morphology of dendritic spines, and were filtered according to custom parameters, those verified with a cluster size distribution analysis of each protein (GluR1, Rab11a, LAMP1). Cluster sizes were determined by setting visual thresholds for each stain individually for 5 images, followed by fitting a Gaussian curve to the obtained distribution to calculate mean puncta sizes. Spots were determined to be inside the dendrite (dendritic spots) using the Spots Close to Filament Border IMARIS extension, with distances for each protein signal determined based on the cluster size distribution analysis (0.18 μm for GluR1, Rab11a and LAMP1). Surfaces were manually filtered according to the dendritic spots (dendritic surfaces).

#### IMARIS analysis: Dendrite

After dendrites were digitally reconstructed, dendrite information was exported as csv files containing values for the dendrite area, position, volume, spine length, spine area, spine volume, and spine position.

#### Characterization of dendritic surfaces

Dendritic surfaces, as described in the reconstruction section, were first exported as a batch. Following this, the same surfaces were manually assigned as a spine head or neck-associated puncta (protein clusters) and then their parameters were exported on a per-spine basis. Each surface export resulted in csv files containing values for the surface area, volume, intensity, position, and percentage of overlap with neighboring surfaces.

#### Colocalization of dendritic spots

Dendritic spots were determined to be colocalized using the IMARIS colocalize spots extension, using a distance of 0.25 μm based on the cluster size distribution analysis.

## Dendritic spine confocal analysis

To determine dendritic spine density images were acquired using an inverted laser scanning confocal microscope (Nikon Instruments, Nikon Eclipse Ti with C2Si+ confocal) controlled via NisElements (Nikon Instruments, version 4.51) using a 488 nm laser. Images were taken with a 60x Plan Apo objective (Nikon Instruments; Plan Apo, NA 1.4, oil immersion objective) and z-stack images were acquired using a step size of 0.2 μm. The brightest dendrites were selected from each image to ensure that the dim protrusions, such as filopodia, were reliably identified. To reliably identify spines of a given dendrite, we analyzed GFP-positive dendrites that were not occluded by other nearby cell processes. Dendrite selection and all following spine analyses were conducted blindly. Dendritic spine density was calculated using the Filament reconstruction function in IMARIS. Each target dendrite was first manually traced using the filament tool. The center of the trace was set to automatic to allow the software to best estimate the shape of the dendrite. Each dendritic filament was approximately 50-70 μm in length. Only unbranched, single stretches of dendrites ending in tufts were used for spine density analysis. Following dendrite tracing using the filament tool, the IMARIS slice view function was used to estimate the thinnest spine heads as well as largest spine lengths (0.15 and 3.10 μm, respectively). These numbers were then used to reconstruct spines, allowing for branching (this refers to branched spines being allowed as a reconstruction, and counted as separate spines). Next, each spine was manually assessed for accuracy using position of spine terminal points and spines were either deleted or drawn in as was necessary. Once we obtained the number of spines per dendritic branch, we divided this number by the total dendritic branch length (> 50 μm) to obtain spine density values.

## Electrophysiological recordings and analysis

Mice (P20-25) were anesthetized with a 4% isoflurane-oxygen mixture (v/v) and perfused intracardially with an ice-cold Perfusion/Slicing artificial cerebrospinal fluid (P/S aCSF) bubbled with 95% O_2_/ 5% CO_2_ containing the following (in mM): 3 KCl, 26 NaHCO_3_, 1.25 NaH_2_PO_4_, 212 sucrose, 10 glucose, 0.2 CaCl_2_, and 7 MgCl_2_ (300-310mOsm). Coronal slices (300 μm thickness) were cut in the same P/S aCSF using a VT1000 S (Leica, Wetzlar, Germany) vibratome before being transferred to a Recording aCSF solution bubbled with 95% O_2_/ 5% CO_2_ containing the following (in mM): 125 NaCl, 2.5 KCl, 25 NaHCO_3_, 1.4 NaH_2_PO_4_, 16 glucose, 0.4 Na-ascorbate, 2 Na-pyruvate, 2 CaCl_2_, and 1 MgCl_2_ (300-310mOsm). Slices were incubated in this Recording aCSF for 30 min at 35 °C before being allowed to recover at room temperature for 1 h prior to recording. All recordings were performed at 29-31°C. Only electroporated neurons in L2/3 of the prelimbic, infralimbic, and anterior cingulate cortex divisions of the mPFC were recorded from. Signals were recorded with a 5X gain, low-pass filtered at 6 kHz, and digitized at 10 kHz using a patch-clamp amplifier (Multiclamp 700B, Molecular Devices, San Jose, CA).

Whole-cell voltage-clamp recordings were made using 3-5 MΩ borosilicate pipettes filled with an internal solution that contained the following (in mM): 120 Cs-methane sulfonate, 8 NaCl, 10 HEPES, 10 CsCl, 10 Na_2_-phosphocreatine, 3 QX-314-Cl, 2 Mg^2+^-ATP, 0.2 EGTA (293 mOsm, adjusted to pH 7.3 with CsOH). Series resistance (Rs) and input resistance (Rin) were monitored throughout the experiment by measuring the capacitive transient and steady-state deflection in response to a −5 mV test pulse, respectively. For mEPSCs recordings, cells were voltage clamped at −70 mV in the presence of tetrodotoxin (TTX, 1μM, Tocris, Bristol, United Kingdom) and picrotoxin (PTX, 100 μM, HelloBio). Liquid junction potentials were left uncompensated. mEPSCs were identified and their amplitude, frequency, rise, and decay determined using custom scripts written in MATLAB (MathWorks, Natick, MA). At least 120 s were analyzed for each cell. Cells were excluded if Rs varied by more than 5 MΩ or exceeded 20 MΩ at any point during the recordings. For each condition, 20-25 cells were recorded from a total of 4-6 animals across 2-3 litters. All electrophysiology reagents were purchased from Sigma-Aldrich (St. Louis, MO) unless otherwise stated.

## Statistical analysis

For confocal image analysis and electrophysiological recordings, we focused on L2/3 neurons in the anterior cingulate cortex, prelimbic, infralimbic, and medial orbital divisions of the mPFC. All statistical analysis was completed in GraphPad Prism 8.0 (GraphPad Software, San Diego, CA) and the threshold for significance for all tests was set to 0.05 (α = 0.05). Spine density and electrophysiological data were analyzed using a one-way ANOVA followed by Tukey’s posttest or a student’s t test. Both male and female mice were used, and we have previously shown that increased levels of C4 in L2/3 mPFC pyramidal cells do not cause sex-dependent differences in connectivity or PFC-associated behavior in mice^8^. Analysis was performed blind to condition. Figures were prepared using CorelDRAW Graphics Suite X8 (Corel Corporation, Alludo, Ottawa, Canada) and ImageJ (NIH, Bethesda, MD). Data are presented as the Mean ± SEM, unless otherwise noted.

## DATA AVAILABILITY

Data are available at https://osf.io/7em3s/?view_only=0e7ffde4ebd344dc83af83b5a605c451

## CODE AVAILABILITY

Custom-written routines are available at https://github.com/CruzMartinLab/IMARIS_compiler and https://github.com/CruzMartinLab/Ephys_analysis_code.

## Extended Data Figures

**Extended Data Figure 1:**
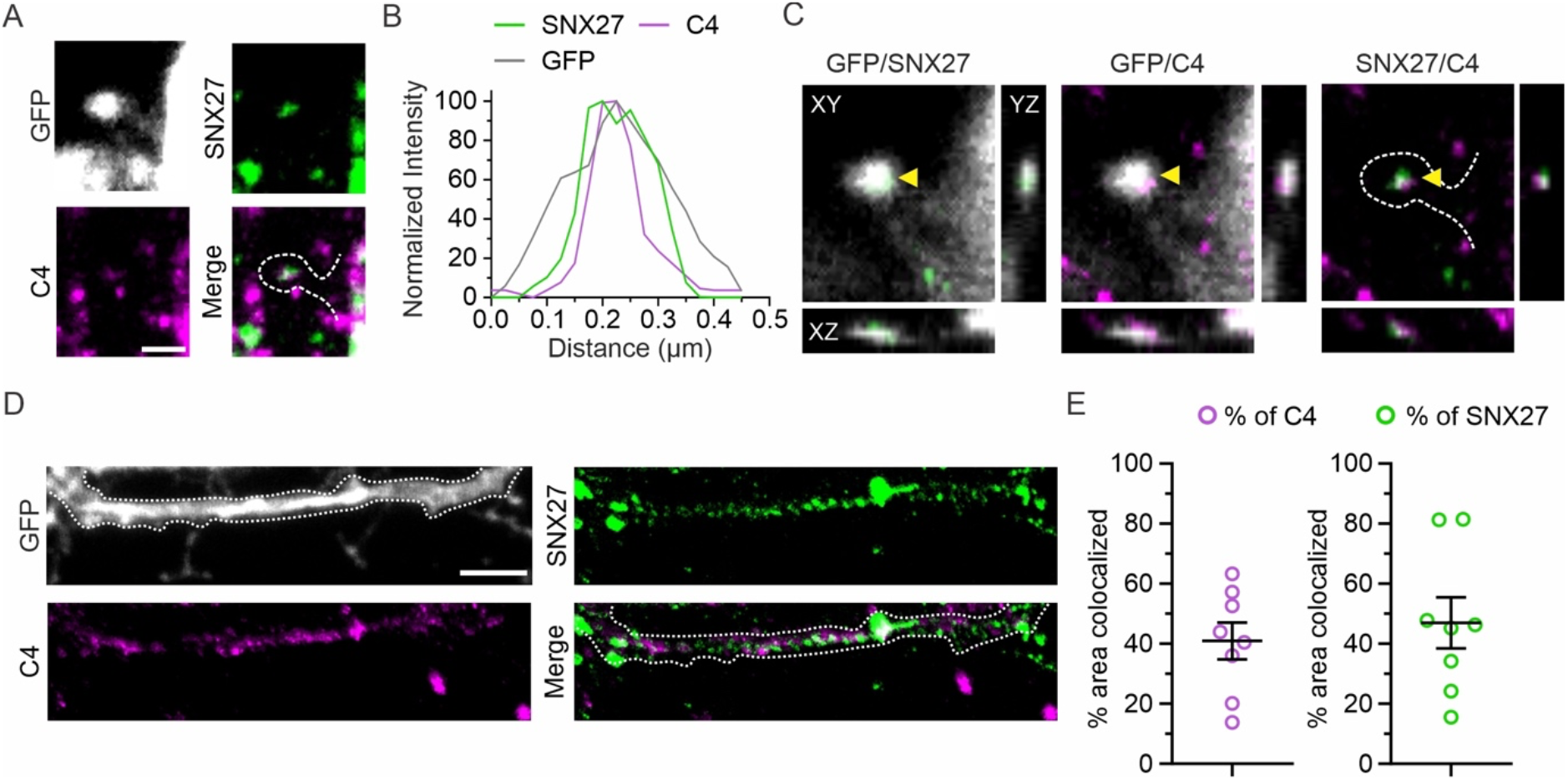
Super-resolution imaging reveals that overexpressed C4 and SNX27 colocalize in dendrites in cultured prefrontal cortex neurons. (**A**) Representative images (60X) showing GFP-positive dendritic spine (white), SNX27 (green) and C4 (magenta) of DIV 14 cultured prefrontal cortical neurons. White dotted line indicates spine morphology. Scale bar = 0.5 μm. (**B**) Normalized intensity profile of a line drawn through a spine head and colocalizing SNX27 and C4 clusters in panel (**A**) showing intensities of GFP (grey line), SNX27 (green) and C4 (magenta). (**C**) Orthogonal views of panel (**A**) are shown (XY, YZ and XZ). Yellow arrowhead indicates an instance of SNX27 and C4 colocalization in the spine. (**D**) Representative images (60X) showing GFP-positive dendrites (white), SNX27 (green) and C4 (magenta) of DIV 14 cultured prefrontal cortical neurons, were used for colocalization analysis. All channels of the image were individually thresholded to calculate area colocalization of C4 and SNX27 in the dendrite. White dotted line indicates dendritic morphology. Scale bar = 2 μm. (**E**) Analysis of cluster signals of overexpressed C4 and SNX27 showed that approximately 41% of C4 area colocalized with SNX27 and approximately 44% of SNX27 area colocalized with C4 in dendrites. *N*=7 dendritic segments, from 3 cells. All graphs, Mean ± SEM.

**Extended Data Figure 2:**
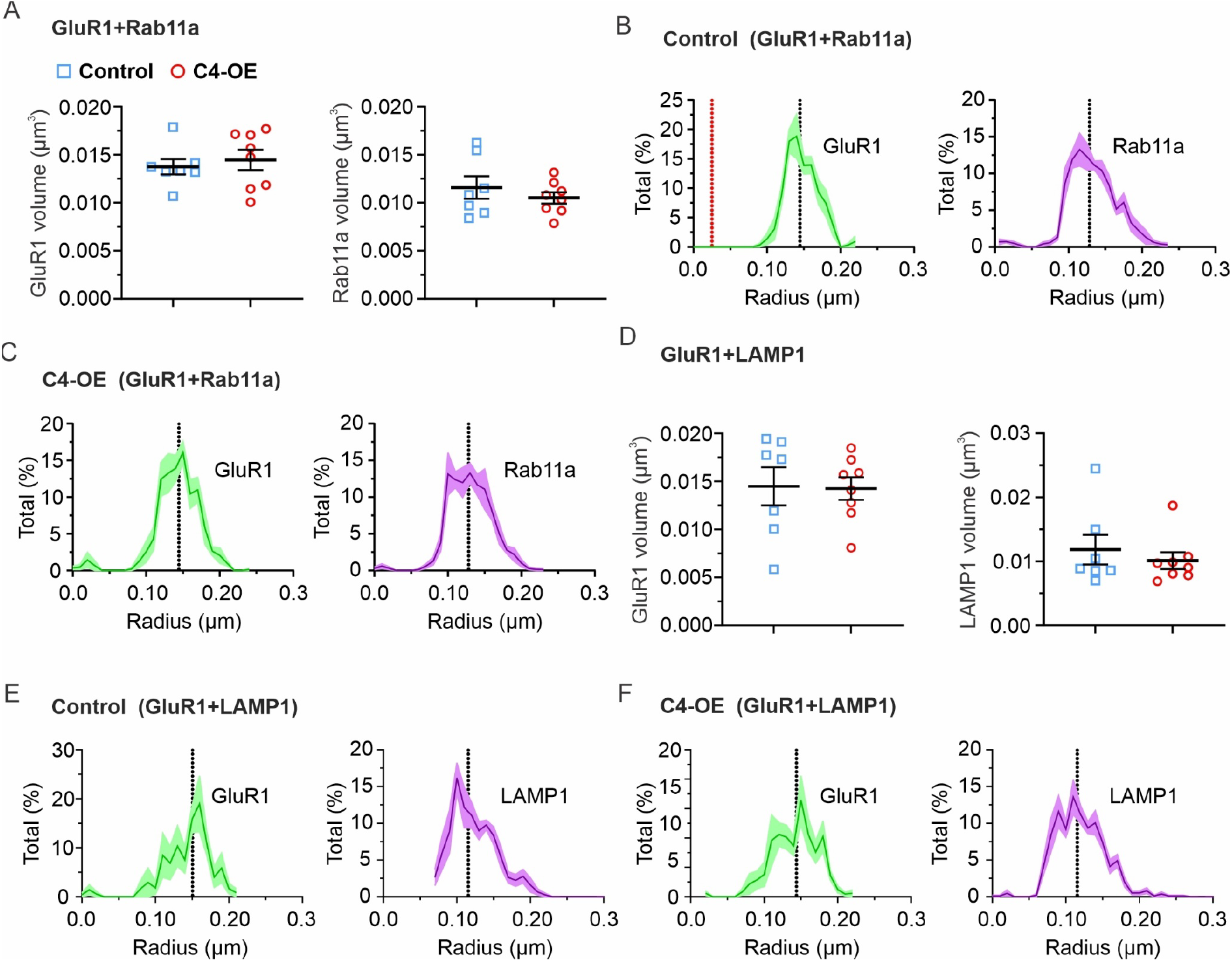
Super-resolution imaging reveals no change in protein cluster sizes of GluR1, Rab11a and LAMP1 with C4-OE. (**A**) C4-OE did not lead to any change in average cluster volume of GluR1 (left panel) or Rab11a (right panel). (**B, C**) Cluster radii distribution of GluR1 (green) and Rab11a (magenta) in GFP-control (**B**) and C4-OE (**C**) respectively. (**D**) C4-OE did not lead to any change in average cluster volume of GluR1 (left panel) or LAMP1 (right panel). (**E, F**) Cluster radius distribution of GluR1 (green) and LAMP1 (magenta) in GFP-control (**E**) and C4-OE (**F**) respectively. (**A-C**) *N*=8 dendrites, 3 animals for GFP-control and C4-OE. (**D-F**) *N*=7 dendrites, 3 animals for GFP-control and *N*=8 dendrites, 3 animals C4-OE. GluR1 was co-stained in the same brain slice with Rab11a (**GluR1+Rab11a**) or with LAMP1 (**GluR1+LAMP1**). All statistics are with student’s t test. All graphs, Mean ± SEM.

